# Hyperconnectivity of the default mode network in multiorgan dysfunction syndrome

**DOI:** 10.1101/418160

**Authors:** Antonio Jimenez-Marin, Diego Rivera, Victoria Boado, Ibai Diez, Fermin Labayen, Irati Garrido, Daniela Ramos-Usuga, Javier Rasero, Alberto Cabrera, Sebastiano Stramaglia, Juan Carlos Arango-Lasprilla, Jesus M. Cortes

## Abstract

Multiple Organ Dysfunction Syndrome (MODS) is a systemic physiological disorder affecting two or more body organs triggered after an insult complication. Beyond the systemic failure, patients who survive MODS present cognitive and neurological impairments that remain stable even several years after Intensive Care Unit (ICU) discharge. Here, we focus on the specific situation of MODS patients with no apparent brain damage (NABD), where the mechanisms driving cognitive impairment at long term are not well-understood. We recruit N_1_ = 13 MODS patients with NABD at 6 months after ICU discharge, together with N_2_ = 13 healthy controls (matched by age, sex and years of education), and acquire functional magnetic resonance imaging at rest to find that, as compared to control, MODS patients with NABD present an overall increase of the functional connectivity (FC) at rest. In particular, we find that the default mode network (DMN) hyperconnects (increasing the node strength of the FC matrix) to three classes of networks: primary sensory (such as auditory, sensory-motor and visual), multimodal integration (such as dorsal attention and salience) and higher order cognition networks (such as fronto-parietal, language and executive control). Therefore, although these patients do not have an apparent structural damage after MODS, at the functional level, we found brain network alterations coexisting with hyperconnectivity of the DMN, that similar to what happens at the onset of other pathologies, might indicate a possible mechanism for brain compensation occurring after MODS.

## Introduction

Multiple Organ Dysfunction Syndrome (MODS), also known as multiorganic failure, is a physiological systemic disorder affecting two or more body organs triggered after a life-threatening insult [1]. Historically, the treatment and care of MODS patients by physicians in the Intensive Care Unit (ICU) has been focused on patient’s survival. However, accumulated evidence have shown that current clinical practice is not optimal, as patients who survive MODS have prevalent sequelae affecting multiple brain-related domains, such as cognitive (attention, executive function, speed processing), neurological (fine psychomotricity, coordinated motor skills, sleep problems) and psychiatric levels (anxiety, depression, post-traumatic stress disorder), alterations that, although potentially reversible, can last even for several years after ICU discharge [2, 3, 4, 5].

With regard to the amount of the cognitive impairment following MODS, the authors in [6] showed that the global cognition level of MODS patients (as measured by the Repeatable Battery for the Assessment of Neuropsychological Status, RBANS) was lower than the one for Mild Cognitive Impairment patients even one year after the insult, and this occurred independently on patient’s age, although for patients older than 65 years, the cognitive impairment after MODS arrived even to the level of the Alzheimer’s disease.

Some possible mechanisms might explain the severe cognitive impairment occurring on MODS patients right after the insult. Independently on the etiology, either an infectious, toxic or traumatic insult triggers a systemic inflammatory response, releasing some of the proinflammatory cytokines such as IL1, IL6, IL8 or TFN who activate the immune system, producing a proliferation and infiltration of lymphocytes and histiocytes in several organs for a defensive response [7, 8]. But sometimes, the inflammatory response becomes pathological, in a manner that the normal immune defensive role converts into deleterious, producing endothelial damage, microvascular dysfunction or tissue oxygenation alterations that can affect the central nervous system, causing for instance, encephalopathies, deficit of consciousness, brain hemodynamic alterations (such as arterial hypotension and hypoperfusion shock) or respiratory insufficiency (arterial hypoxemia) [9, 10]. To which extent these mechanisms can produce cognitive and neurological alterations at long term is not well-understood.

In some cases, even with no apparent brain damage (NABD) –meaning that no brain lesions are detected by a neuroradiologist from structural imaging–, whether or not MODS patients after ICU discharge present immune system’s hyperactivation that can propagate to the CNS is still unknown, and the research regarding to the cognitive impairment in these patients is almost inexistent. Indeed, it is of crucial interest to understand the pathophysiological mechanisms of these brain functioning aftermaths, in order to somehow improve patient’s quality life in a short-time [11].

Clinical evidence has shown some variables have an association with cognitive impairment in these patients after ICU discharge, such as advanced age, hypoxemia, hypotension, the amount of sedation days and the amount of delirium days [12, 13, 14, 6]. Little is known about neuroimaging biomarkers in these patients [15].

Here, we hypothesize that functional neuroimaging can differentiate between MODS patients with NABD and healthy participants. In particular, we focus on brain’s functional organization at the large scale by studying the functional connectivity (FC) at rest [16, 17, 18], a condition occurring when the brain participant is not involved in any particular task. During resting, different brain connectivity techniques such as seed-based correlation analysis, independent component analysis or partial least squares decomposition can separate brain dynamics into a variety of resting state networks (RSNs) [19, 20, 21, 22, 23, 24], namely, patterns of synchronous dynamics that resemble different activation maps when performing specific tasks, such as for instance, auditory, visual, sensory-motor, executive-control or the default mode network (DMN) [25].

Of special interest for clinical studies, it has been widely shown that different RSNs become altered in different pathological conditions such as deficit of consciousness [26, 27, 28, 29, 30], schizophrenia [31, 32], epilepsy [33], Alzheimer’s Disease [34, 35, 36, 37, 38, 39] and healthy aging [40]. As far as we know, what RSN is altered after MODS has not been yet addressed.

## Materials and Methods

### Ethical considerations

The study was approved by the local Ethical Committee in the Cruces University Hospital (Code CEIC E16/52, IPs: Jesus M Cortes and Juan Carlos Arango-Lasprilla).

MODS patients were treated in the Cruces University Hospital and recruited at the moment of ICU discharge by the physician patient’s responsible, who collected the information consents from the patients.

Healthy controls (HC) were recruited from patient’s representatives, who provided different information consents as patients, only valid for HC, as required by the ethics committee.

### Participants

The study included a number of N= 26 participants, N_1_ = 13 MODS patients and N_2_ = 13 HC. Both groups were matched by age (MODS: 51.54 ± 9.36 year; HC: 52.92 ± 10.17 years; p-value= 0.72 after two-sample t-test), sex (MODS: 8 males; HC: 6 males; p-value= 0.43 after chi-squared test) and years of education (MODS: 13.46 ± 5.46; HC: 13.61 ± 5.22; p-value= 0.94 after two-sample t-test). Demographic data for both MODS and HC are given respectively in Tables 1 and 2.

**Table 1:** Demographic data for MODS patients. M = male, F = female, N/A = information not available, RI = Respiratory insufficiency, SS = Septic shock, AES = Acute endocarditis surgery, HS = Hypovolemic Shock

| ID | Age | Gender | Education level | SOFA | Cause of injury |
| --- | --- | --- | --- | --- | --- |
| P01 | 31 | F | 13 | 4 | RI |
| P02 | 40 | M | 20 | 6 | RI |
| P03 | 46 | F | 21 | 4 | SS |
| P04 | 46 | F | 21 | 5 | HS, RI |
| P05 | 46 | M | 9 | 10 | SS |
| P06 | 52 | F | 8 | 6 | SS |
| P07 | 54 | M | 9 | 10 | RI |
| P08 | 55 | M | 22 | 8 | SS |
| P09 | 57 | M | 12 | 5 | RI |
| P10 | 59 | F | 10 | 14 | AES |
| P11 | 59 | M | 8 | 6 | SS |
| P12 | 61 | M | 10 | 7 | RI |
| P13 | 64 | M | 12 | 8 | SS |

**Table 2:** Demographic data for Healthy Controls.

| ID | Age | Gender | Education level |
| --- | --- | --- | --- |
| **C01** | 29 | M | 8 |
| **C02** | 45 | M | 21 |
| **C03** | 45 | M | 18 |
| **C04** | 45 | F | 5 |
| **C05** | 50 | F | 15 |
| **C06** | 54 | M | 7 |
| **C07** | 55 | F | 18 |
| **C08** | 56 | F | 18 |
| **C09** | 56 | F | 16 |
| **C10** | 60 | F | 8 |
| **C11** | 62 | F | 18 |
| **C12** | 65 | M | 11 |
| **C13** | 66 | M | 14 |

Patient’s inclusion criteria was to have a SOFA score ≥ 4. The SOFA clinical scale, acronym of Sequential Organ Failure Assessment, roughly measures how many organs *fails* after the insult. For an exact definition, see Table 1 in the reference [41]. Notice that MODS is really a severe pathology, as SOFA ≥ 4 has an associated mortality rate of at least 20% [42], and can be higher, eg., 60% for SOFA between 8 and 11.

Patient’s exclusion criteria was to have a score in the Mini–Mental State Examination (MMSE) *<* 23, any magnetic resonance imaging contraindication, visible abnormalities in brain structural imaging (T1 sequence), having had cerebral hipoxia during MODS, had a history before MODS of surgical intervention in CNS, drug abuse, ophthalmological, neurological, psychiatric or cardiovascular diseases, potentially influencing imaging or clinical measures.

HC’s exclusion criteria was a score in the Mini–Mental State Examination (MMSE) *<* 23, a score in the depression questionnaire PHQ-9*>* 4, magnetic resonance imaging contraindications, visible abnormalities in brain structural imaging (T1 sequence), had a history of surgical intervention in CNS, drug abuse, ophthalmological, neurological, psychiatric or cardiovascular diseases, potentially influencing imaging or clinical measures.

### Imaging acquisition

Magnetic resonance imaging (MRI) was performed with a Philips 3-tesla Achieva Dstream MRI scanner with a 32-channel head coil. Imaging acquisition was performed in both MODS and HC six months after recruitment, as we were interested in cognitive functioning at long term.

#### Anatomical data

High resolution T1 images were acquired with a 3D Turbo Field Echo (TFE): repetition time TR = 7.4 ms, echo time TE = 3.4 ms, voxel size = 1.1 × 1.1 × 1.2 mm^3^, slice thickness = 1.2 mm, field of view FOV = 250 × 250 mm^2^, 300 contiguous sagittal slices covering the entire brain and brainstem.

#### Resting state functional data

were acquired with a total duration of 7.40 minutes and using the following parameters: 214 whole-brain gradient echo echo-planar images with TR/TE = 2100*/*27 ms, FOV = 240 × 240 mm^2^, voxel size = 3 × 3 × 3 mm^3^, 80 × 80 matrix, slice thickness of 3 mm, 45 axial slices, interleaved in ascending order.

### Imaging preprocessing

We applied a preprocessing for the resting-state fMRI similar to that used in previous work [43, 44, 45, 46, 47, 48], by using FSL and the FEAT toolkit [49, 50]. First, a MCFLIRT is performed in order to correct the head movement artifacts; next, slice-time correction was applied to the fMRI dataset. After intensity normalization, the functional data were spatially normalized to the MNI152 brain template, with a voxel size of 2 × 2 × 2 mm^3^. Finally, we regressed out the motion time courses, the average cerebrospinal fluid signal and the average withe-matter signal. All voxels were then spatially smoothed with a 6 mm full-width-at-half-maximum isotropic Gaussian kernel and a bandpass filter was applied between 0.01 and 0.08 Hz [51], followed by the removal of linear and quadratic trends.

### Craddock’s Functional Atlas

We made use of a functional atlas of 2754 regions of interest (ROIs) following the unsupervised clustering method published in [52]. After providing 3000 as an input (equal to the number of desired ROIs), the algorithm spatially constrained the different voxels belonging to the same region to be spatially contiguous whilst maximized both within-region similarity and between-region difference. From the initial 3000 desired regions, the algorithm only provided 2754 valid ROIs, who were the ones satisfying the algorithm constrains.

### Mask for the resting state networks

Following [25], we created a mask for different RSNs by defining voxels with z-score value satisfying *z <* −3 or *z >* 3. In particular, we built masks for the default mode, cerebellum, executive control, frontoparietal, sensorimotor, auditory and visual resting state networks.

These masks were used for two different approaches: 1. To calculate strength brain maps of connectivity from all ROIs to each specific RSN and, 2. To calculate the overlapping between the brain maps resulting from group comparison and each different RSN.

We also made use of the following networks: visual, sensory-motor, auditory, fronto-parietal, executive control, language, dorsal attention (lateral visual) and ventral attention (salience), obtained from single-subject in [53] and available at https://findlab.stanford.edu/functional_ROIs.html.

### Brain maps of functional connectivity strength

After averaging voxel time series within a given ROI, each subject was represented with 2754 time series, each with T= 214 time points. Per subject, we obtained one FC matrix of dimension 2754 × 2754 by calculating pairwise Pearson correlation for all pairs of time series. Brain maps of FC strength per subject were obtained by summing over either rows or columns the FC matrix.

In addition to the strength maps obtained from the all-to-all FC matrix, i.e., when using the 2754 × 2754 FC matrix to build the strength maps, we also calculated strength maps from alternative FC matrices built by calculating the correlations from all ROIs in the brain to the ROIs belonging to each specific RSN mask, thus obtaining rectangular (rather than square) FC matrices of dimension 2754 × (RSN Size). The strength maps were calculated by summing in the rectangular FC matrix over the dimension of the specific RSN. In particular, we made use of the following FC matrices: all-to-DMN,all-to-cerebellum, all-to-executive control, all-to-frontoparietal, all-to-sensorimotor, all-to-auditory and all-to-visual matrices.

### Statistical analysis

Group comparison, MODS vs HC, was performed using the *randomise* function available in FSL using the images of strength maps, one per subject and divided in the two groups. Brain maps of group differences were obtained with two different contrasts, [1 -1] (patients *>* control) and [-1 1] (patients *<* control).

Statistical significance was approached by permutation testing using the method of Threshold-Free Cluster Enhancement (TFCE) implemented in *randomise*, a non-parametric method for finding significant clusters after group comparison without a priory defining any cluster or region of interest. For further details see https://fsl.fmrib.ox.ac.uk/fsl/fslwiki/Randomise/UserGuide.

### Separability of group matrices measured by the signal to noise ratio (SNR)

From signal-detection theory, two important parameters control the differences between two groups (here, MODS and HC). On one side, the mean of the two distributions should be the more different (ie., separated) as possible. On the other, the two distributions should be the more sharp (small standard deviation) as possible. Indeed, these two factors can be combined by defining the signal to noise ratio 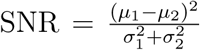, where *μ*_*i*_ and 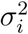 represent mean and variance in group *i* (for further details see [54]). Therefore, the higher the SNR, the bigger separability of the two distributions.

## Results

N= 26 subjects participated in the study. N_1_ = 13 MODS patients and N_2_ = 13 HC were recruited in the vicinity of the Cruces University Hospital (Bilbao, Spain). Structural and functional magnetic resonance imaging were acquired for each participant.

We have studied a specific subgroup of MODS patients having no apparent brain damage (NABD). Figure 1 shows three axial slices illustrating ventricle space, together with white and gray matter for three randomly chosen MODS patients. Even for a neuroradiologist, it is hard to see differences between the three images as compared to HC. More quantitative neuroimaging methods, such as voxel-based morphometry as implemented in FSL did not provide any significance group differences. Therefore, both qualitative and quantitative standard methods for analyzing structural imaging did not show any structural damage in MODS patients with NABD.

**Figure 1:**
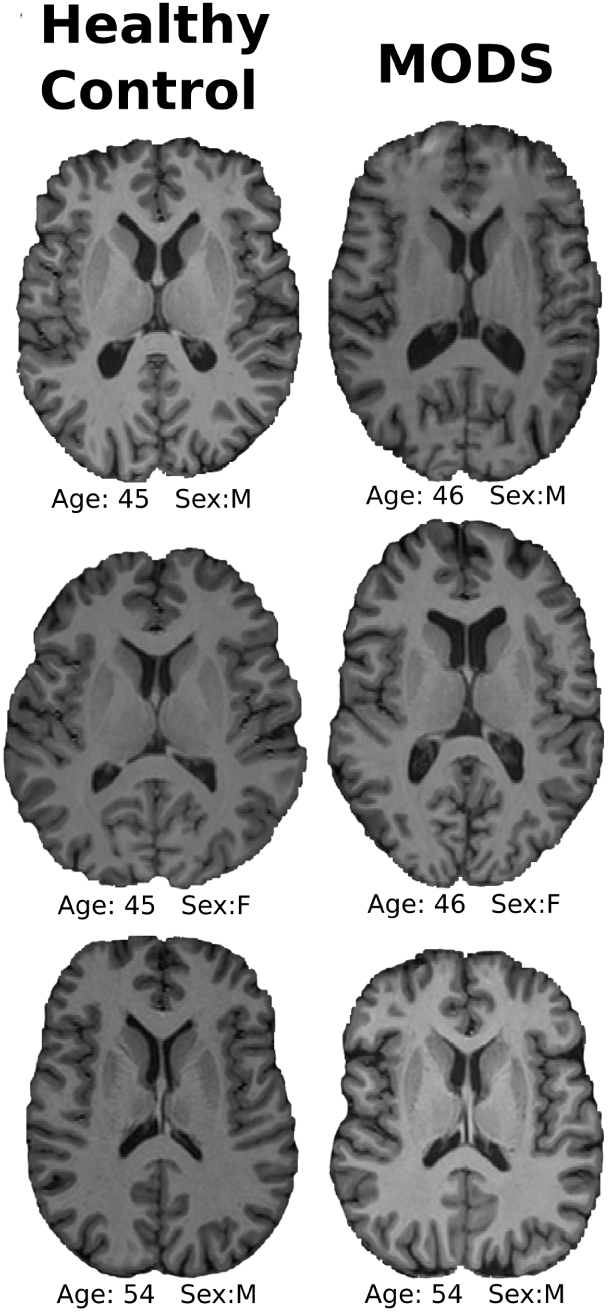
MODS patients with no apparent brain damage (NABD). Even for a neuroradiologist, it is hard to find differences between MODS patients and HC. Here, we illustrate three axial slices of the T1 image for both HC and MODS. Age and sex are indicated on the bottom of each image.

Next, we asked whether group differences existed by analyzing functional neuroimaging data. We first studied the association between brain maps of strength and the clinical SOFA scale (methods), and no significant regions survived after multiple comparison.

Second, we searched for group differences in brain maps of strength (methods). No significant differences were found for the strength maps obtained from the all-to-all FC matrix, that with dimension 2754 × 2754 was built calculating the Pearson correlation between the time series from all to all ROIs.

Third, also to perform group comparison, we calculated the strength from other matrices built by calculating the correlations from all ROIs in the brain to the ROIs belonging to each specific RSN mask (methods). Figure 2 illustrates the method for the all-to-DMN FC matrix, but the same analysis was repeated for the all-to-cerebellum, all-to-executive control, all-to-frontoparietal, all-to-sensorimotor, all-to-auditory and all-to-visual matrices.

**Figure 2:**
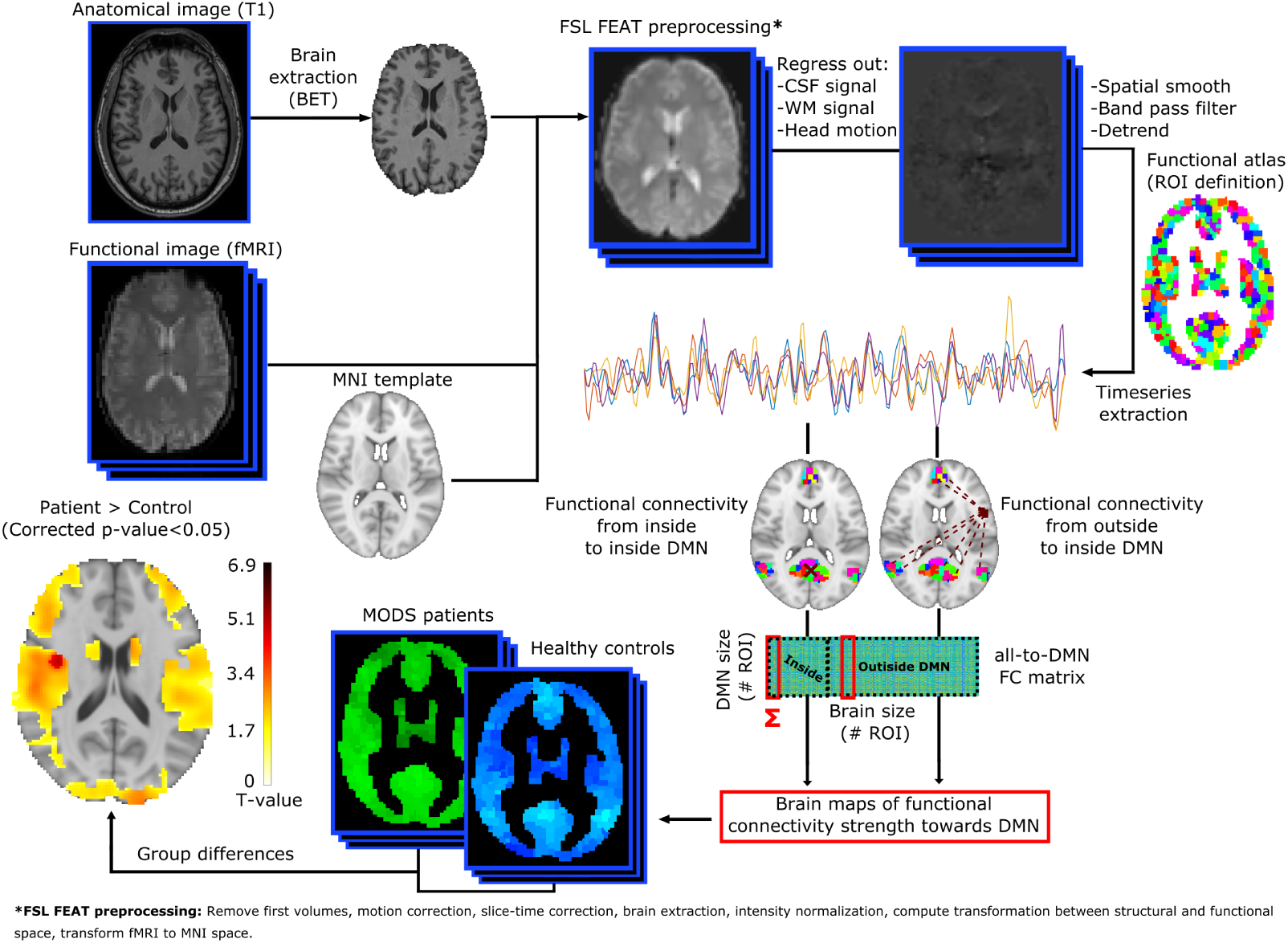
General pipeline for the analyzing of the functional data at rest. Double acquisitions were acquired, consisting in high-resolution anatomical images (T1) and functional images at rest. Following state-of-the-art image preprocessing, time-series of the blood oxygenation level dependent signal were obtained for each region of interest (ROI), defined by a functional atlas of 2754 ROIs. Different functional connectivity (FC) matrices have been calculated, all-to-all (accounting for the correlations from all ROIs to all ROIs), all-to-DMN (from all ROIs to only those ROIs belonging to the DMN, illustrated here in this figure), all-to-cerebellum, all-to-executive control, all-to-frontoparietal, all-to-sensorimotor, all-to-auditory and all-to-visual. Summing over the dimension of the RSN, we calculated strength brain maps, each per subject, and performed group comparison correcting for multiple comparison. For details see methods.

The variability of the strength measure depended on RSN size (figure 3a), thus applying this strategy for small RSNs we were capable of controlling intra-group variability. Figure 3b illustrates the probability distribution of strength values calculated from the all-to-all FC matrix for the two groups: HC (colored in blue) and MODS (in red).

**Figure 3:**
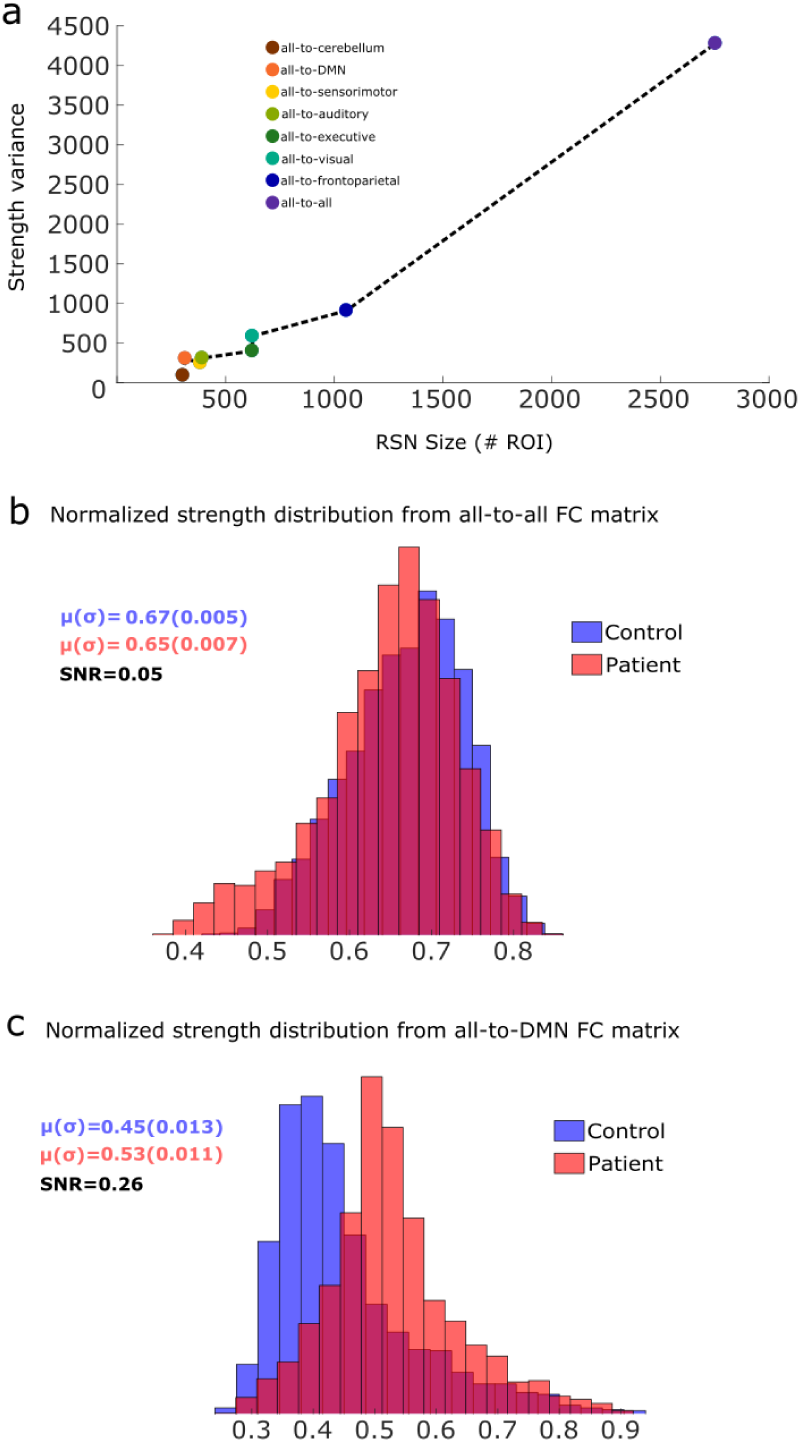
Controlling intra-group variability by calculating strength maps over different FC matrices. **a:** Variability, defined as the variance of the strength values, from different classes of FC matrices as a function of the size (number of regions of each RSN). Here, strength values for control and patients have been pooled together. In general, variability scales with size. **b:** Probability distributions for the two groups, control (blue) and patients (red), of the strength obtained from the all-to-all FC matrix. Mean and standard deviation, together with the signal to noise ratio (SNR) are provided for the two groups. **c:** Similar to b, but now the strength distributions have been calculated using the all-to-DMN FC matrix. Notice that the separability of the two distributions, as measured by the SNR, have increased from 0.05 in panel b to 0.26 in panel c, an increment ratio of 520% by using the all-to-DMN rather than the all-to-all FC matrix. In addition, patient’s distribution shifted to the right, indicating hyperconnectivity. For both panels a and b, dark red areas represent the intersection of the two distributions.

Figure 3c illustrates the distribution of strength values from the all-to-DMN FC matrix. In comparison to the all-to-all FC matrix, the separability of the two distributions increased a factor of 520%, as the SNR was equal to 0.05 for the all-to-all and 0.26 for the all-to-DMN (for details on SNR, see methods). Values of SNR for all other RSNs are smaller than the one for all-to-DMN (see all values in Table 3).

**Table 3:** Signal to Noise Ratio (SNR) obtained from different types of FC matrices, each associated to a different resting state network. In all cases, SNR is obtained from the strength distribution in HC and MODS groups. The first two values of the table correspond to the distributions depicted in figure 3, all-to-all in panel 3b and all-to-DMN in panel 3c.

| Type of FC matrix: | SNR |
| --- | --- |
| all-to-all | 0.049 |
| all-to-DMN | 0.26 |
| all-to-Sensorimotor | 0.051 |
| all-to-auditory | 0.015 |
| all-to-visual | 0.018 |
| all-to-frontoparietal | 0.00034 |
| all-to-cerebellum | 0.014 |

After group comparison, the contrast [-1 1] (patients *<* control) did not provide any significant differences after multiple comparison in any class of the FC matrix. As a consequence, no significant hypoconnection were found in any of the RSNs.

For the contrast [1 -1] (patients *>* control) the only situation which provided significant group differences occurred for the all-to-DMN, and no other FC matrix did it (figure 4). Importantly, when the SNR was calculated only over regions belonging to these maps of hyperconnectivity, we got an increase of SNR from 0.26 (figure 3c) to 0.63 (figure S1), indicating that the higher significance in the group differences, the higher SNR.

**Figure 4:**
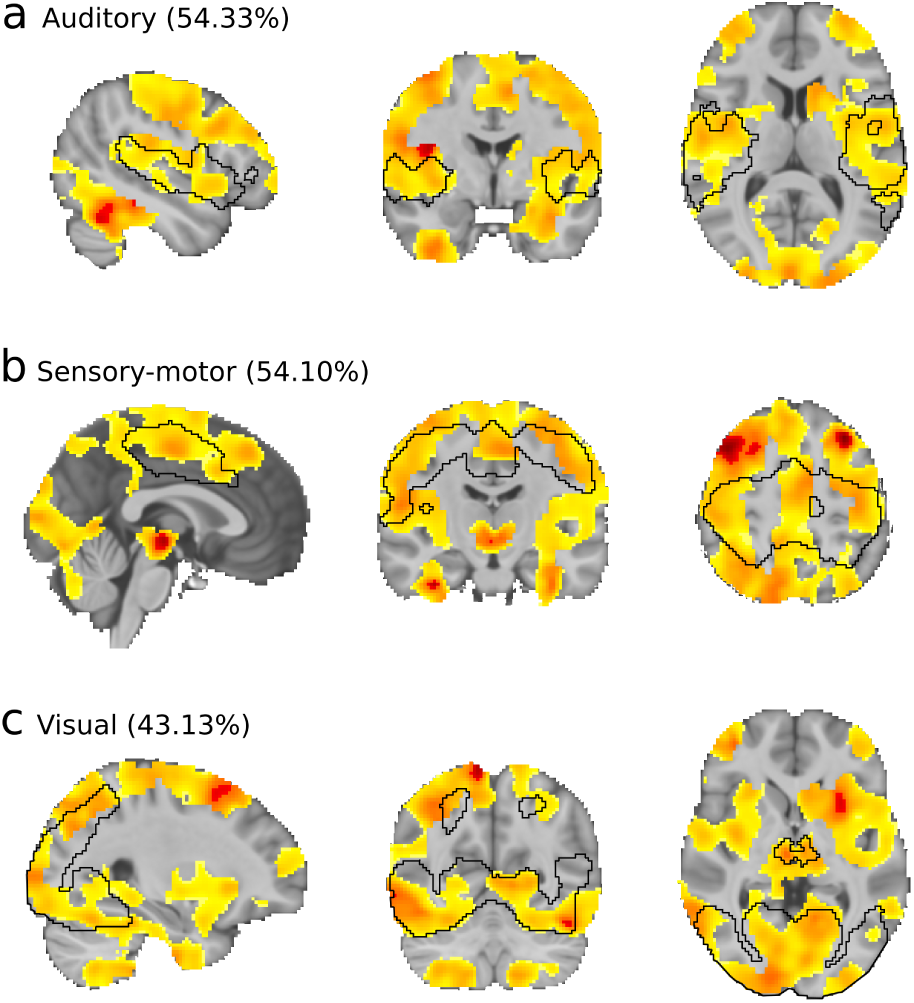
MODS patients show hyperconnectivity of the DMN majorly to auditory, sensorimotor and visual networks. **a-c:** Brain maps resulting from group comparison after multiple corrections of the strength maps with the contrast [1 -1], therefore, patient *>* control connectivity. **a:** In black, we depict the mask of the auditory RSN. Brain maps after group comparison overlapped 54.33% with the auditory network. Similar information is provided in panels b and c, respectively, for the (**b**) sensorimotor and (**c**) visual networks.

Hyperconnectivity, here identified by the increase of strength values in MODS as compared to HC, was found only in the DMN and majorly towards three RSNs: auditory (matching with the mask 54.33%), sensorymotor (matching 54.10%) and visual (matching 43.13%). Anatomically, areas of hyperconnection from the DMN to the auditory RSN were bilaterally found in Heschl’s gyri, rolandic operculum, temporal superior and insula. To the sensorimotor network, we found bilaterally the precentral and postcentral gyri, supperior motor area, middle cingulum and inferior parietal. Finally, hyperconnection to the visual network were bilaterally found in inferior, middle and superior occipital cortices, lingual gyrus, cerebellum, calcarine sulcus, fusiform, temporal inferior cortex and cuneus. Notice that, from the functional data analysis we cannot say these brain structures have any structural damage but that the oxygenation dynamics in these areas are more hyper-connected to the DMN than in HC.

## Discussion

Multiple organ dysfunction syndrome (MODS) is a progressive disorder triggered by a life-threatening insult that independently on the etiology of the injury has an associated high mortality rate [42]. Albeit MODS has not been yet classified as a neurological disorder, as ICU physicians have been traditionally focused on patient’s survival, however, approximately half of the patients who overcome MODS evince post-intensive care sequelae [55], that having an impact on psychological, neurological and psychiatric levels [2, 3, 4, 5] can last even for several years after the MODS insult [6]. Despite on the importance of improving patient’s intervention during ICU for a better outcome, little is known about the neural mechanisms of the brain-related alterations at long term in MODS patients.

Here, we have analyzed FC at rest in MODS patients six months after ICU discharge, and found hyperconnectivity of the DMN in these patients, that as far as we know, has not yet been reported. Of crucial importance, hyperconnectivity of the DMN has been suggested before as a network plasticity mechanism for brain-damage compensation, occurring in the onset of a plethora of pathologies, for instance, after concussion [56], in the early stage of Alzheimer’s disease [57], in patients who survive to brain tumors [58], deficit of consciousness [59, 60], cancer related cognitive impairment [61], autism [62] and behavioural disorders such as anorexia [63]. Therefore, hyperconnectivity of the DMN is not specific to MODS.

We have shown that the brain maps of DMN hyperconnectivity affected majorly to primary sensory networks such as auditory, sensory-motor and visual (figure 5). However, the brain maps of DMN hyperconnectivity also overlapped with multimodal integration networks [64], mainly with the dorsal and ventral attention (a.k.a. salience) networks, indicating that the hyperconnectivity of the DMN in MODS patients also affect to intermediate stages in the information processing from sensory to higher order cognitive networks (such as fronto-parietal, executive control and language), which provide further evidence supporting the compensation strategy for brain recovery after MODS.

**Figure 5:**
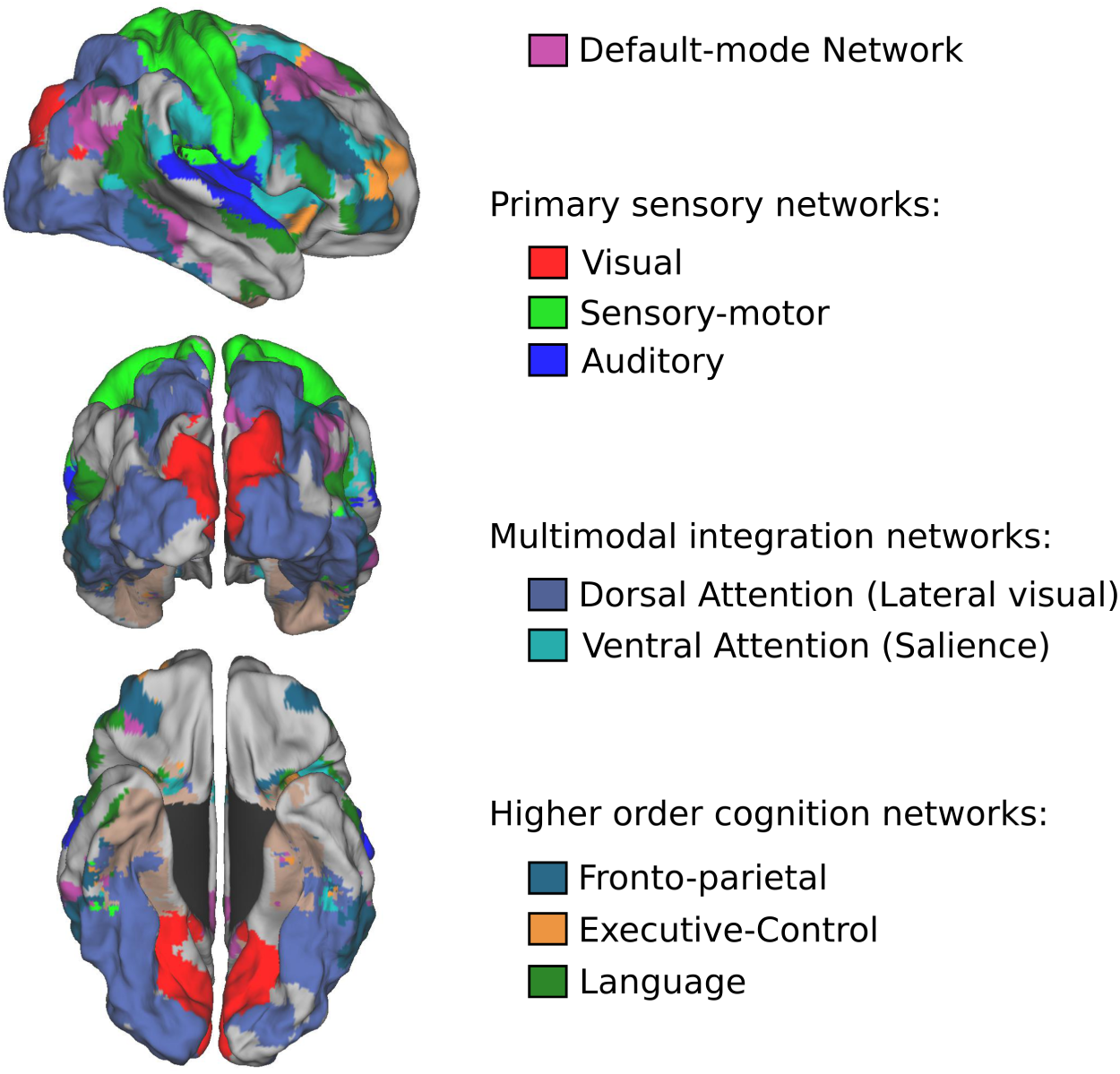
Hyperconnectivity of the DMN might indicate a compensation strategy for brain damage after MODS. The DMN (here colored in pink) hyperconnects to primary sensory networks (visual, sensory-motor and auditory), to multimodal integration networks (dorsal and ventral attention) and to higher order cognitive networks (fronto-parietal, executive control and language).

Future studies combining clinical and physiological variables during ICU, together with neuroimaging and neuropsychological evaluations are needed to fully characterize neural mechanisms underlying cognitive impairment in MODS patients.

## Acknowlegments

ID acknowledges financial support from Department of Education of the Basque Country, postdoctoral program. JR acknowledges financial support from the Minister of Education, Language Policy and Culture (Basque Government) under Doctoral Research Staff Improvement Programme. We acknowledge financial support from Department of Economical Development and Infrastructure of the Basque Country, Elkartek Program (grant no. KK-2018/00032), Ministerio Economia, Industria y Competitividad, Spain and FEDER (grant no. DPI2016-79874-R) and Fundacion Mutua Madrileña. JMC and JCAL are funded by Ikerbasque.

---

**Figure S1:**
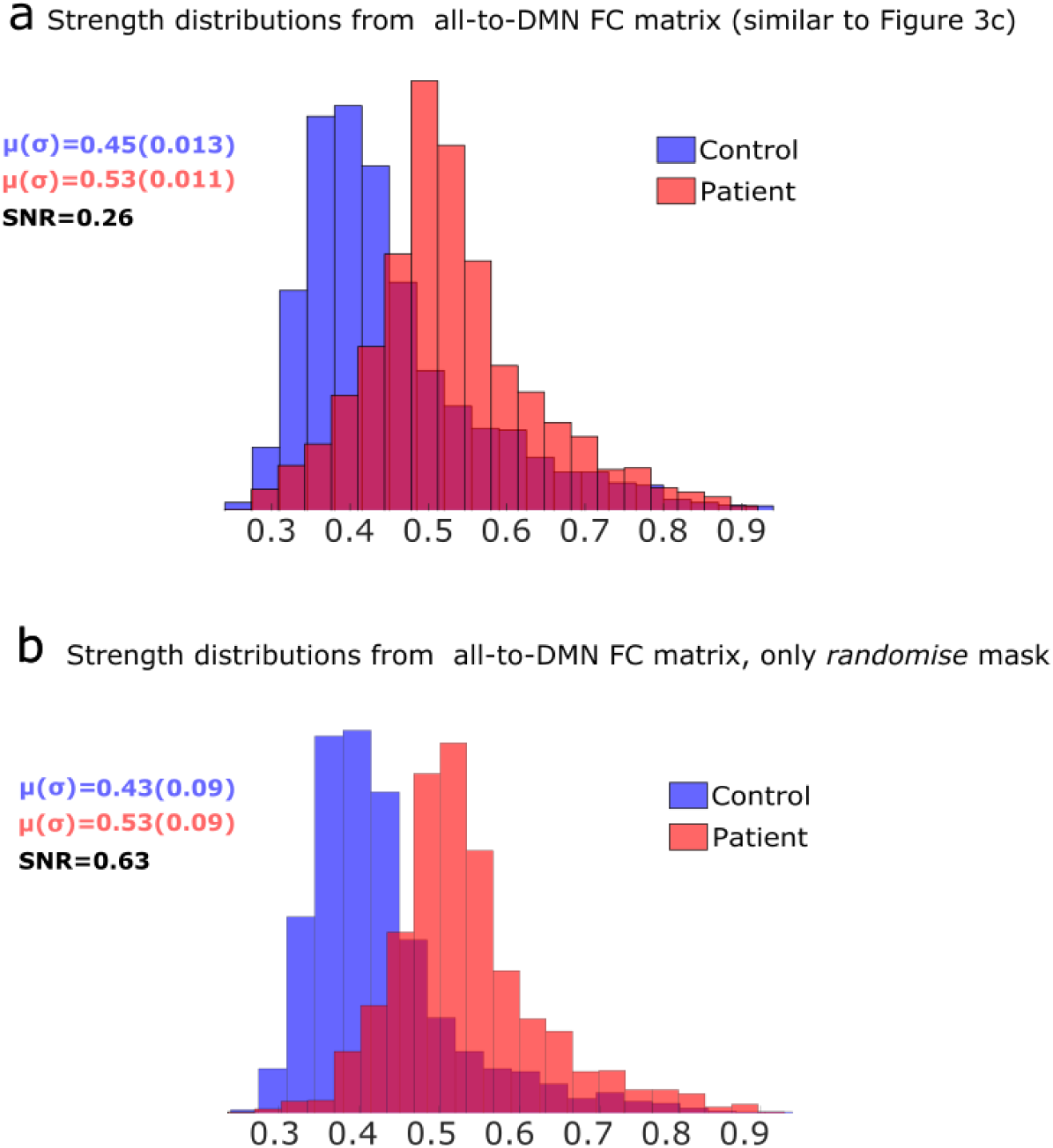
All-to-DMN strength distributions for control and patients. **a:** Same plot as in figure 3c, replicated here for illustration purposes. **b:** Strength distributions for both control and patients but calculated only on the ROIs that intersect with the maps of significant differences after group comparison (using the *randomise* function implemented in FSL). Notice that SNR can increase from 0.26 in panel a to 0.63 in panel b, therefore increasing a factor of 242% in these specific regions.

